# Escobase and AcR: *Escherichia coli* databases of genomes and acid resistance genes from isolates from cattle farms

**DOI:** 10.1101/2025.01.12.632593

**Authors:** Martin Lucile, Ruiz Philippe, Leroy Sabine, Sapountzis Panagiotis

## Abstract

*Escherichia coli* is a gram-negative bacterium commonly found in mammals and birds’ intestinal microbiota. It can be easily isolated and has been studied for more than nine decades, making it an ideal model organism. It is mainly commensal but occasionally pathogenic, expressing virulence factors, antibiotic-resistance and acid-resistance genes, which contribute to its ability to persist and colonize diverse environments. Its multi-resistance, combined with the virulence of certain pathotypes (e.g. STEC), poses a recurring threat to public health, as treatment for clinical infections is not always straightforward. The remarkable propensity of *E. coli* for diversification and genomic variability is often referred to as an open pangenome.

This project’s objective was to create the ‘Escobase’ database (**Es**cherichia **co**li data**base**), which is comprised exclusively of *Escherichia coli* isolates from cattle farms, allowing us to generate a reference set of genomes that can be used for comparative genomics on samples from cattle and cattle farms. Using the annotated *E. coli* genomes, we further built a catalog of predicted proteins that help *E. coli* survive in low pH environments, often encountered when the bacteria traverse through the intestine. We believe these two databases will be of great help to the community, and we plan to keep improving them and publish updated versions in the future.

## Introduction

*Escherichia coli* is a gram-negative gamma-proteobacterium (1) present in the environment and the GastroIntestinal Tract (GIT) of many animals, including *Bos taurus*. While most *E. coli* strains are commensal, the species is exceptionally diverse and also encompasses a range of pathotypes. Cattle is a major reservoir of Shiga Toxin *Escherichia coli* (2), a.k.a. STEC, which is a pathotype that presents a significant health risk for humans. Other pathogenic *E. coli* strains include the enteropathogenic *E. coli* (EPEC), the enterotoxigenic *E. coli* (ETEC), the enteroinvasive *E. coli* (EIEC), the enteroagggregative *E. coli* (EAEC), the diffusely adherent *E. coli* (DAEC) and the invasive adherent *E. coli* (AIEC) (2). The virulence of these strains is variable, from very low (and thus essentially avirulent, which is often the case in cattle) to hypervirulent, causing life-threatening conditions in humans. Even though the total number of cases per year is not very high, there are no viable treatment options for several *E. coli* pathotypes, making them one of the most lethal pathogens, especially for the most sensitive age groups (pre-school children and elderly).

*E. coli*’s ability to colonize and infect is often aided by virulence genes (3, 4), which can be disseminated across strains via mobile genetic elements such as plasmids and bacteriophages, and are often present in pathogenicity islands (DNA segments carrying virulence factors).

Antibiotic Resistance Genes (ARGs) encode functions that render bacteria tolerant to antibiotics. They are frequently found in *E. coli* isolated from cattle feces, and since various classes of antibiotics are commonly used in cattle farms, there is an increased pressure for the emergence of various AMR (Anti Microbial Resistance) bacterial genes, which comprises the resistome of the digestive tract (5).

Acid resistance genes (AcR), which have been studied less (compared to the AMR and virulence genes), are genes that are expressed in response to stress in an acidic environment, such as compartments of the digestive tract (6). Common strategies to tolerate increased acidity include the modification of the membrane and the porins that constitute it, allowing a reduction in the influx of protons into the cell, and changes in the chaperone proteins that limit the damage caused by the acidic environment. These genetic elements have likely contributed to *E. coli*’s ability to successfully colonize the GIT of various hosts, including humans.

This project, which was part of a MSc thesis at the University of Clermont Auvergne (UCA) in Clermont-Ferrand (France), aimed at building an *E. coli* genome database comprised exclusively of *E. coli* isolates from bovine feces. Our *E. coli* database (Escobase) can be used as a reference dataset when comparing sequence data (from genomes and metagenomes) from cattle farms. The data used originate from several studies, including the HECTOR project by Ferrandis-Vila et al. (7) and Leekitcharoenphon et al. (8), while the rest of the data come from *E. coli* genomes that were available on NCBI in May 2023. As part of this project, the predicted aminoacid sequences of the *E. coli* genomes were compared to a Virulence Factor database (VFdb), an AMR database (ResFinder), and a custom-made Acid Resistance database (AcR) from predicted aminoacid sequences that likely help the bacteria survive the low pH encountered in the GIT of humans and animals.

## Methods

Handling of files, data storage and bioinformatic analyses were carried out on the computing cluster of the Mésocentre of the University of Clermont Auvergne, and locally in a linux/bash and Conda v23.3.1 environment (Conda contributors, https://docs.conda.io/projects/conda/). All tables were compiled, edited, and exported in R (4.4.2) and Rstudio (2024.04.2) using the packages ‘dplyr’,’tidyr’ and ‘tidyverse’ (9). Tables were further edited in Libreoffice 24.2.7.2. Heatmaps were generated in R using the ‘heatmap’ function.

### Datasets Recovery and Processing

A total of 915 *E. coli* genomes were used for this project, which originated from the HECTOR project, the Leekitcharoenphon et al. project. (2021; PRJEB41365) (7, 8), and additional studies that were available at the NCBI (National Center for Biotechnology Information) in May 2023 (Table ‘[Metadata-Annot-Escobase_lm20230710_refined.csv](https://entrepot.recherche.data.gouv.fr/file.xhtml?fileId=176228&version=DRAFT)’). We arrived at this number by retaining only the *E. coli* genomes from isolates originating from cow feces (out of the total of *E. coli* genomes that were available in May 2023), and which included additional information such as the place/country of origin, the date of collection, and the methodology used to produce the genomes. Data handling was performed with GNU bash v4.4.20(1)-release, and data retrieval was carried out with the Entrez-direct v16.2 tool (10) from NCBI, and tables were examined and edited on LibreOfficeCalc v6.0.7. Additional information on the genomes, such as the N50, genome size, their number of CDS, and the ATCG ratio, were retrieved with the QUAST v5.0.2 tool.

### Genome assembly and annotation

For the aforementioned data, we prioritized downloading the genome assemblies and annotation data when they were available. If there was no assembly, we downloaded the sequencing reads (exclusively Illumina reads) and assembled them using SPAdes v3.15.0 (11). Contigs were then submitted to Prodigal v2.6.3 (12) to identify open reading frames and predict coding sequences and proteins.

### Characterization of AMR, virulence and acid resistance genes

Antibiotic resistance (ARGdb), virulence factor (VFdb) and a custom-made acid resistance (AcRdb) database, were either downloaded or built for this study. For the acid resistance database, we first created a set of genes, which have been identified by previous studies as key factors providing acid resistance in *E. coli* strains in low pH environments (Table 1), and then searched for these genes in the downloaded *E. coli* genomes that were already assembled and annotated (n=886, NCBI annotation pipeline). We used cd-hit (13) to create a non-redundant dataset of the predicted aminoacid sequences (eliminating only sequences that were 100% identical and the alignment covered the 100% of their length) and used it as a reference protein database (501 aminoacid sequences from 886 genomes). The full dataset of proteins of the Virulence factor database (14) was downloaded by the www.mgc.ac.cn/VFs/ website in July of 2022, and the ResFinder dataset of AMR genes was downloaded on the first of October of 2022. For each of the three protein datasets, we built a diamond blast database using the ‘makedb’ command with default settings (Diamond v2.0.5; 15). All Diamond blast comparisons were performed using an Evalue of 1e-50, a minimum percentage identity of 75%, and a minimum subject and query coverage of 70% as cutoffs.

**Table 1:**
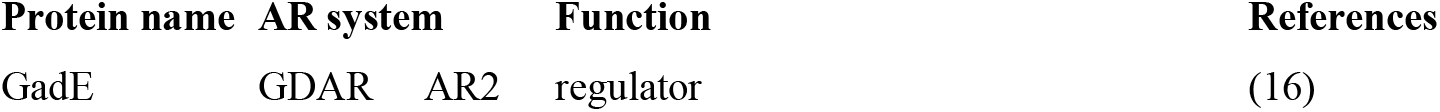
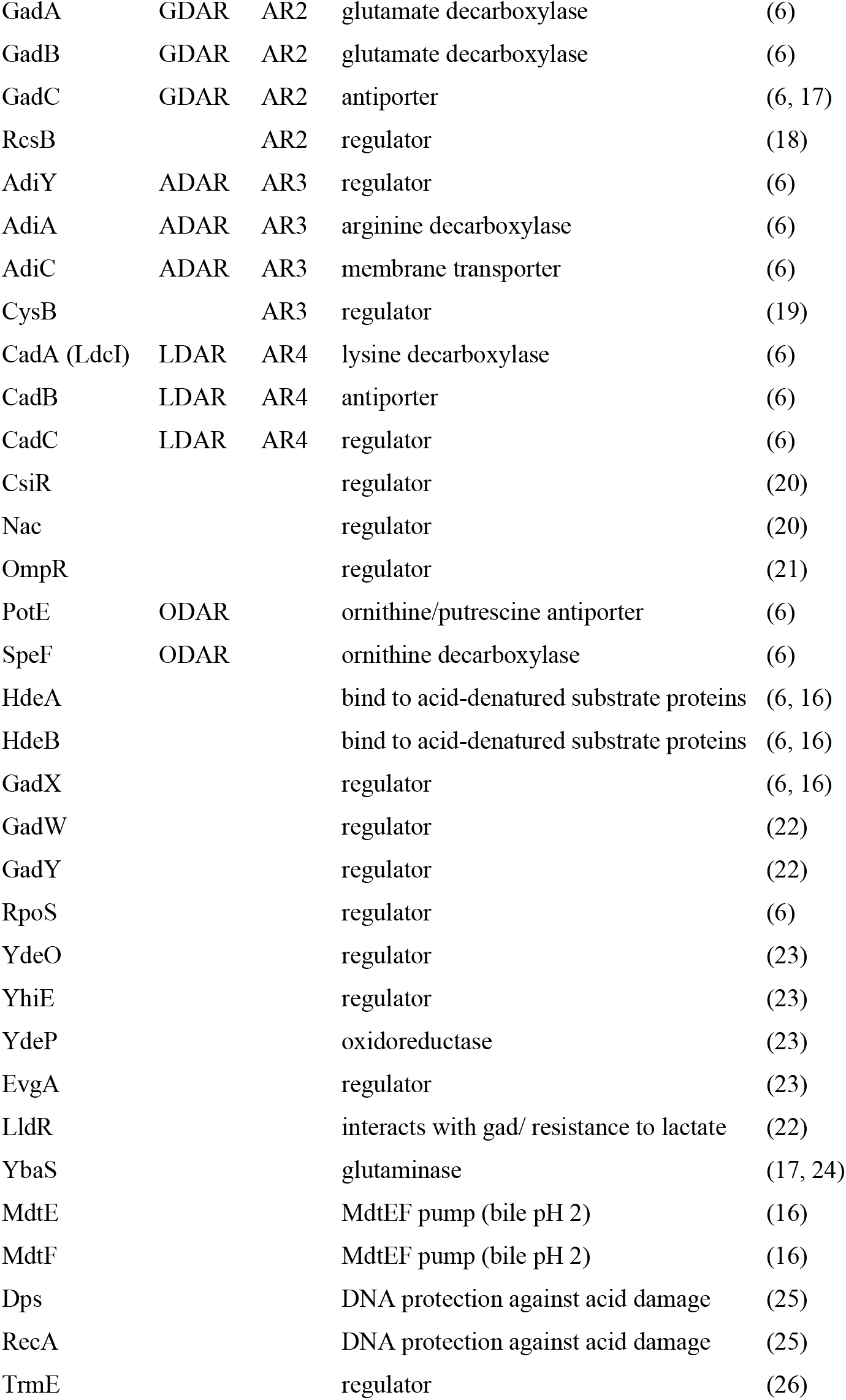
Acid resistance genes selected for the AcR database. From left to right: the protein name, the acid resistance system to which it belongs, and the function it serves, as well as the study it came from.

| Protein name | AR system | Function | References |
| --- | --- | --- | --- |
| GadE | GDAR AR2 | regulator | (16) |

|  |  |  |  |  |
| --- | --- | --- | --- | --- |
| GadA | GDAR | AR2 | glutamate decarboxylase | (6) |
| GadB | GDAR | AR2 | glutamate decarboxylase | (6) |
| GadC | GDAR | AR2 | antiporter | (6, 17) |
| RcsB |  | AR2 | regulator | (18) |
| AdiY | ADAR | AR3 | regulator | (6) |
| AdiA | ADAR | AR3 | arginine decarboxylase | (6) |
| AdiC | ADAR | AR3 | membrane transporter | (6) |
| CysB |  | AR3 | regulator | (19) |
| CadA (LdcI) | LDAR | AR4 | lysine decarboxylase | (6) |
| CadB | LDAR | AR4 | antiporter | (6) |
| CadC | LDAR | AR4 | regulator | (6) |
| CsiR |  |  | regulator | (20) |
| Nac |  |  | regulator | (20) |
| OmpR |  |  | regulator | (21) |
| PotE | ODAR |  | ornithine/putrescine antiporter | (6) |
| SpeF | ODAR |  | ornithine decarboxylase | (6) |
| HdeA |  |  | bind to acid-denatured substrate proteins | (6, 16) |
| HdeB |  |  | bind to acid-denatured substrate proteins | (6, 16) |
| GadX |  |  | regulator | (6, 16) |
| GadW |  |  | regulator | (22) |
| GadY |  |  | regulator | (22) |
| RpoS |  |  | regulator | (6) |
| YdeO |  |  | regulator | (23) |
| YhiE |  |  | regulator | (23) |
| YdeP |  |  | oxidoreductase | (23) |
| EvgA |  |  | regulator | (23) |
| LldR |  |  | interacts with gad/ resistance to lactate | (22) |
| YbaS |  |  | glutaminase | (17, 24) |
| MdtE |  |  | MdtEF pump (bile pH 2) | (16) |
| MdtF |  |  | MdtEF pump (bile pH 2) | (16) |
| Dps |  |  | DNA protection against acid damage | (25) |
| RecA |  |  | DNA protection against acid damage | (25) |
| TrmE |  |  | regulator | (26) |

## Results

### Overview

The genomes used for the Escobase were selected based on the availability of their metadata, which allowed us to select only the ones isolated from bovine feces, and for which the authors were explicit about their origin, their collection date as well as the context of their study and the sequencing technology used. The Escobase was built using 886 already assembled genomes, and 29 additional ones, which were assembled using the SPADES software (v3.15.0 on the mesocentre cluster), bringing the total to 915 genomes (Table ‘[Metadata-Annot-Escobase_lm20230710_refined.csv](https://entrepot.recherche.data.gouv.fr/file.xhtml?fileId=176228&version=DRAFT)’).

### Functional annotation

For each genome, we first identified the predicted aminoacid sequences, and compared them to reference databases to identify proteins that may be related to virulence traits, or facilitate resistance to antibiotics and an acidic environment. The VFdb and the ResFinder databases were used for the former two comparisons, and for the latter we built an Acid-Resistance (AcR) database using a set of predicted proteins composed of regulators and other proteins implicated as having key roles in acid resistance, based on previous studies (Table 1). By grouping the genomes based on country of origin and presenting their on heatmaps, some variation across the virulence, AMR and AcR features could be observed (Figure 1). However, we refrained from commenting on any of the observed patterns because: 1) the variation is likely linked to multiple factors such as biases due to unequal number of isolates from various locations/countries, the purpose of the study (e.g. exclusively clinical isolates or not), or the variation linked to factors related to animal husbandry practices, diets, antibiotic usage, and 2) such correlations were not the objective of this study.

**Figure 1:**
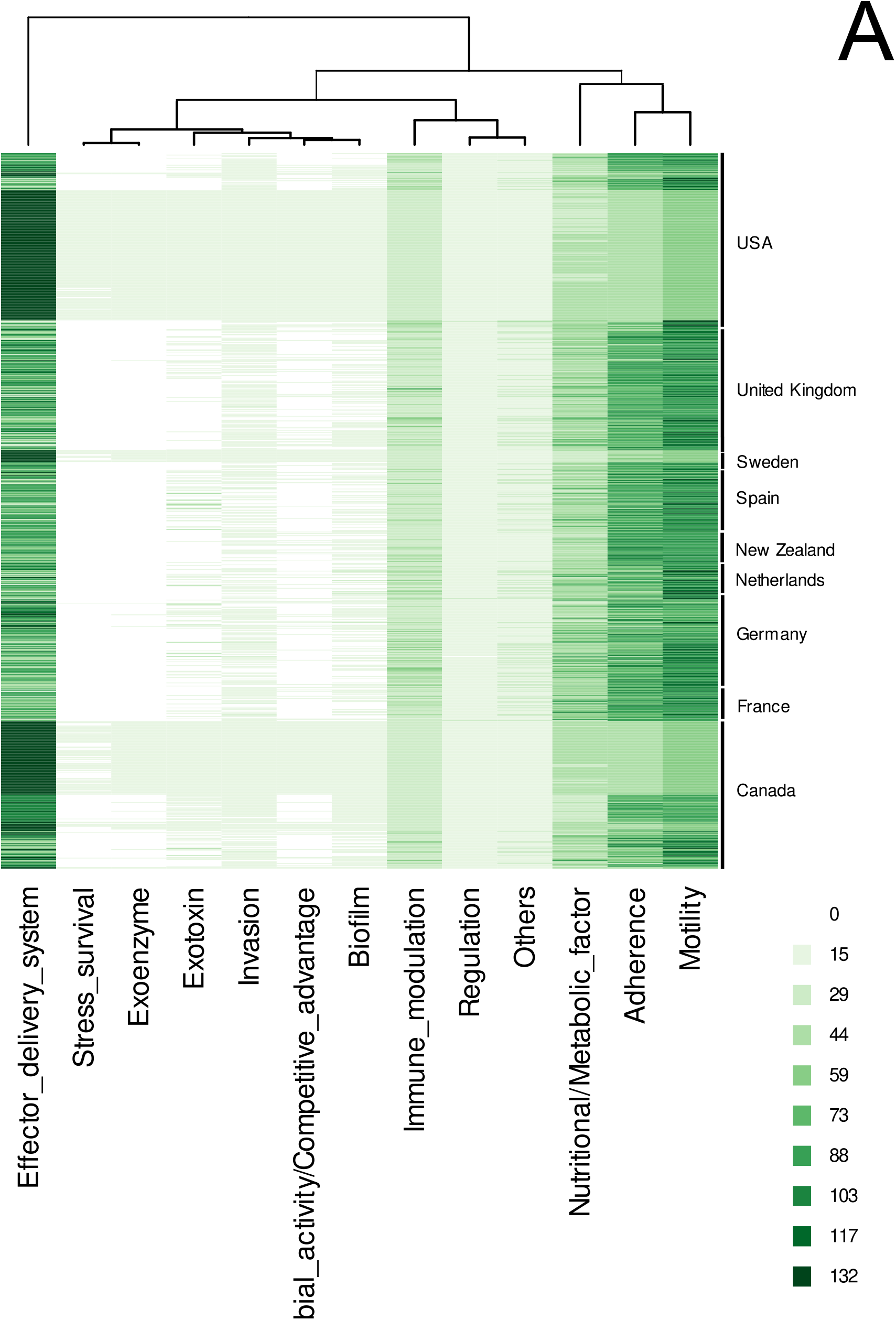
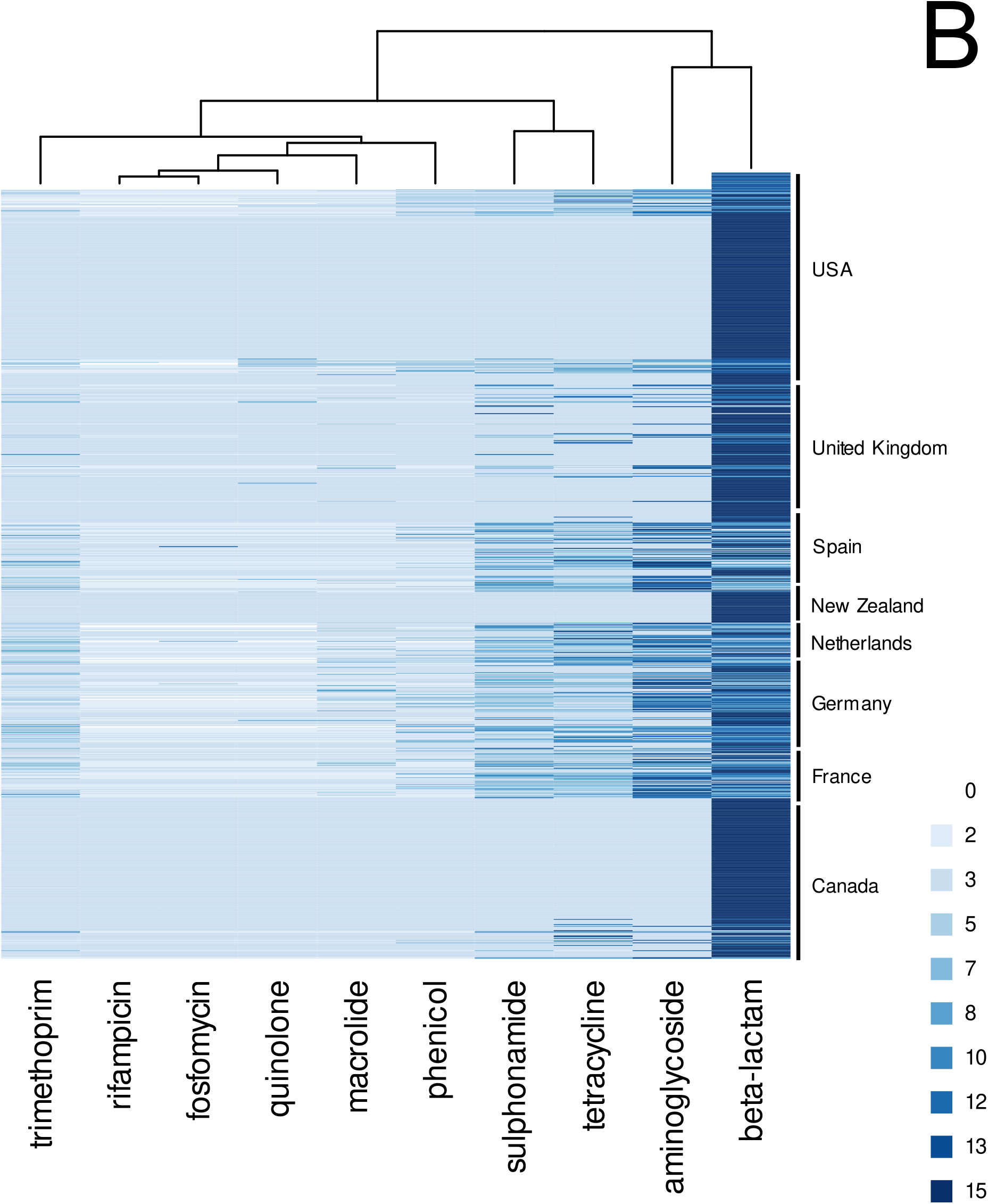
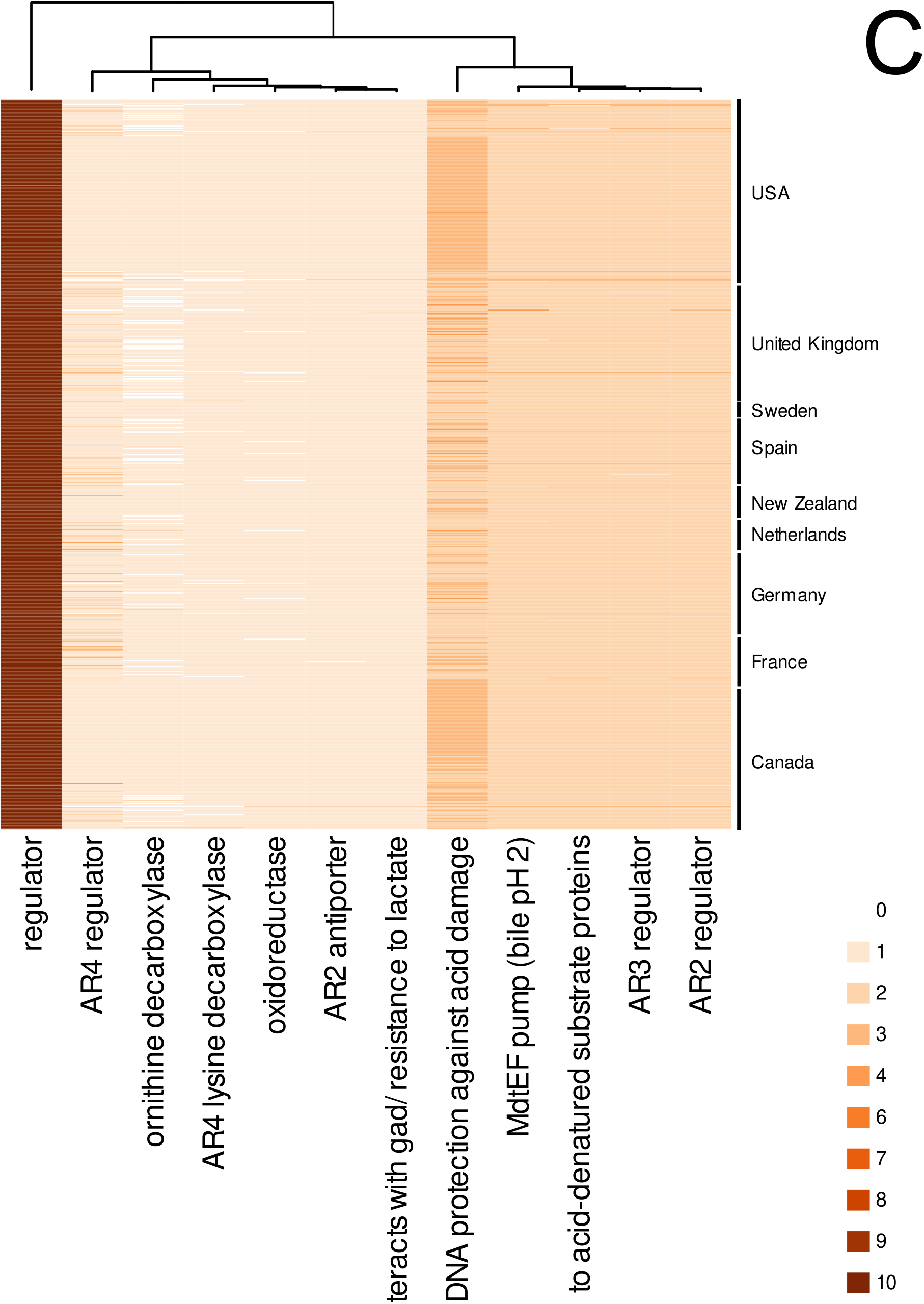
Heatmaps of virulence (A), AMR (B) and acid resistance (C) genes in the 915 genomes of Escobase. Rows show individual genomes and columns the number of genes in each category: A: Virulence category, defined by the authors of VFdb, B: AMR classes, described in the ResFinder database, and C gene functions of the acid resistance genes (Table 1). On top of each heatmap, the hierarchical clustering groups the columns that are more similar based on the distribution of counts. Genomes (rows) have been ordered by country (right).

The most frequently present virulence factors were related to secretion systems (Effector delivery system), followed by factors related to adherence and mobility (Figure 1A). When looking at individual features, the most frequently present in *E. coli* genomes were various Fim-predicted proteins (FimF, FimA, FimG, and FimD; Adherence), AcrB (Antimicrobial activity) and IbeC (Invasion). For the AMR comparisons, we used the ResFinder database, which showed that genes coding for AMR components related to the beta-lactam and aminoglycoside classes were the most frequently present in *E. coli* genomes (Figure 1B), and more specifically the genes encoding for the blaCMY-47_1 and aminoglycoside_aph(3’’)-Ib_2. The most frequently identified genes related to acid tolerance were the ones coding for the transcriptional regulators RcsB, OmpR as well as the RpoS and LidR, but also the DNA starvation/stationary phase protection protein Dps (Figure 1C).

## Discussion and future perspectives

The Escobase and the accompanying AcR database, built for this project, is our first attempt to create a reference genome database of *E. coli* isolates from cattle farms and a gene catalog that can help assess the ability of *E. coli* strains to survive in acidic conditions. These datasets need to be refined and improved both in terms of covering a bigger diversity of genomes (especially considering the open pangenome of *E. coli* species) and genes facilitating acid resistance. This first version of the AcR database includes several predicted ORFs that are regulators and serve multiple functions, including housekeeping that control other physiological functions too (and not only the response to an acidic environment) or component systems that facilitate resistance to acid and several other deleterious compounds (e.g. antibiotics). We chose not to include various genes which have been suggested to facilitate acid resistance because of their wide range of functions (e.g. two component PhoP/PhoQ system), yet we kept in the dataset some housekeeping ones, which were found in all genomes (e.g. RcsB, OmpR regulators), but have been implicated as having key roles in acid resistance (6, 16).

We believe that the Escobase genome database offers a good starting point, which can help researchers to: 1) identify very similar/identical *E. coli* genomes when comparing their own genomes to it, 2) build an MLST (or of a similar principle) system which will use sequence markers (e.g. specific genes) to identify *E. coli* genomes, 3) use it as a reference for phylogenomic, virulence, AMR or any other gene/sequence specific comparisons related to these *E. coli* genomes.

## Data availability

A fasta file containing the 915 genomes of *E. coli* as well as a metatable describing, among others, the date of isolation, origin of the genomes, sequencing technology is available at doi.org/10.57745/RGIZYC. An additional metatable that includes the results of the BLAST output (top hits) using the three aforementioned databases (Vfdb, ResFinder, AcR) is present at the same doi ([Metadata-Annot-colibase_lm20241230_refined_withBLASTs.csv](https://entrepot.recherche.data.gouv.fr/file.xhtml?persistentId=doi:10.57745/IKBKWA&version=1.1)). A fasta file containing the AcR aminoacid fasta file, which was used for building the AcR database, is also available under the same DOI ([AcR_AA_nr_latest30Dec2024.faa](https://entrepot.recherche.data.gouv.fr/file.xhtml?persistentId=doi:10.57745/QZXHLY&version=1.1)). All interested parties are encouraged to contact us for any inquiries, including new additions, suggestions, or edits.

## Notes

https://www.doi.org/10.57745/RGIZYC

### Competing Interest Statement

The authors have declared no competing interest.

## References

1. V. S. Braz, K. Melchior, C. G. Moreira, Escherichia coli as a Multifaceted Pathogenic and Versatile Bacterium. Front. Cell. Infect. Microbiol. 10 (2020).

2. P. Sapountzis, A. Segura, M. Desvaux, E. Forano, An Overview of the Elusive Passenger in the Gastrointestinal Tract of Cattle: The Shiga Toxin Producing Escherichia coli. Microorganisms 8, 877 (2020).

3. H. Sheng, J. Y. Lim, H. J. Knecht, J. Li, C. J. Hovde, Role of Escherichia coli O157:H7 Virulence Factors in Colonization at the Bovine Terminal Rectal Mucosa. Infect. Immun. 74, 4685–4693 (2006).

4. G. Bretschneider, E. M. Berberov, R. A. Moxley, Reduced intestinal colonization of adult beef cattle by Escherichia coli O157:H7 tir deletion and nalidixic-acid-resistant mutants lacking flagellar expression. Vet. Microbiol. 125, 381–386 (2007).

5. D. R. Call, M. A. Davis, A. A. Sawant, Antimicrobial resistance in beef and dairy cattle production. Anim. Health Res. Rev. 9, 159–167 (2008).

6. U. Kanjee, W. A. Houry, Mechanisms of acid resistance in Escherichia coli. Annu. Rev. Microbiol. 67, 65–81 (2013).

7. M. Ferrandis-Vila, et al., Using unique ORFan genes as strain-specific identifiers for Escherichia coli. BMC Microbiol. 22, 135 (2022).

8. P. Leekitcharoenphon, et al., Genomic evolution of antimicrobial resistance in Escherichia coli. Sci. Rep. 11, 15108 (2021).

9. H. Wickham, et al., Welcome to the Tidyverse. J. Open Source Softw. 4, 1686 (2019).

10. J. Kans, “Entrez Direct: E-utilities on the Unix Command Line” in Entrez Programming Utilities Help [Internet], (National Center for Biotechnology Information (US), 2024).

11. A. Bankevich, et al., SPAdes: a new genome assembly algorithm and its applications to single-cell sequencing. J. Comput. Biol. 19, 455–477 (2012).

12. D. Hyatt, et al., Prodigal: prokaryotic gene recognition and translation initiation site identification. BMC Bioinformatics 11, 119 (2010).

13. W. Li, A. Godzik, Cd-hit: a fast program for clustering and comparing large sets of protein or nucleotide sequences. Bioinformatics 22, 1658–1659 (2006).

14. L. Chen, et al., VFDB: a reference database for bacterial virulence factors. Nucleic Acids Res. 33, D325–328 (2005).

15. B. Buchfink, C. Xie, D. H. Huson, Fast and sensitive protein alignment using DIAMOND. Nat. Methods 12, 59–60 (2015).

16. S. H. Schaffner, et al., Extreme Acid Modulates Fitness Trade-Offs of Multidrug Efflux Pumps MdtEF-TolC and AcrAB-TolC in Escherichia coli K-12. Appl. Environ. Microbiol. 87, e00724–21 (2021).

17. P. Lu, et al., L-glutamine provides acid resistance for Escherichia coli through enzymatic release of ammonia. Cell Res. 23, 635–644 (2013).

18. M. D. Johnson, N. A. Burton, B. Gutiérrez, K. Painter, P. A. Lund, RcsB is required for inducible acid resistance in Escherichia coli and acts at gadE-dependent and -independent promoters. J. Bacteriol. 193, 3653–3656 (2011).

19. X. Shi, G. N. Bennett, Effects of rpoA and cysB mutations on acid induction of biodegradative arginine decarboxylase in Escherichia coli. J. Bacteriol. 176, 7017–7023 (1994).

20. P. Aquino, et al., Coordinated regulation of acid resistance in Escherichia coli. BMC Syst. Biol. 11, 1 (2017).

21. A. Stincone, et al., A systems biology approach sheds new light on Escherichia coli acid resistance. Nucleic Acids Res. 39, 7512–7528 (2011).

22. T. Anzai, et al., Expanded roles of lactate-sensing LldR in transcription regulation of the Escherichia coli K-12 genome: lactate utilisation and acid resistance. Microb. Genomics 9, mgen001015 (2023).

23. N. Masuda, G. M. Church, Regulatory network of acid resistance genes in Escherichia coli. Mol. Microbiol. 48, 699–712 (2003).

24. G. Brown, et al., Functional and structural characterization of four glutaminases from Escherichia coli and Bacillus subtilis. Biochemistry 47, 5724–5735 (2008).

25. K. C. Jeong, K. F. Hung, D. J. Baumler, J. J. Byrd, C. W. Kaspar, Acid stress damage of DNA is prevented by Dps binding in Escherichia coliO157:H7. BMC Microbiol. 8, 181 (2008).

26. S. Gong, Z. Ma, J. W. Foster, The Era-like GTPase TrmE conditionally activates gadE and glutamate-dependent acid resistance in Escherichia coli. Mol. Microbiol. 54, 948–961 (2004).

